# Seamless, rapid and accurate analyses of outbreak genomic data using Split K-mer Analysis (SKA)

**DOI:** 10.1101/2024.03.25.586631

**Authors:** Romain Derelle, Johanna von Wachsmann, Tommi Mäklin, Joel Hellewell, Timothy Russell, Ajit Lalvani, Leonid Chindelevitch, Nicholas J. Croucher, Simon R. Harris, John A. Lees

**Affiliations:** NIHR Health Protection Research Unit in Respiratory Infections, National Heart and Lung Institute, Imperial College London, London, W21PG, UK; MRC Centre for Global Infectious Disease Analysis, Department of Infectious Disease Epidemiology, School of Public Health, Imperial College London, London, W12 0BZ, UK; European Molecular Biology Laboratory, European Bioinformatics Institute, Wellcome Genome Campus, Hinxton, UK; Department of Mathematics and Statistics, University of Helsinki, Helsinki 00014, Finland; Centre for Mathematical Modelling of Infectious Diseases, London School of Hygiene & Tropical Medicine, London, UK; Bill and Melinda Gates Foundation, 62 Buckingham Gate, Westminster, London, SW1E 6AJ

## Abstract

Sequence variation observed in populations of pathogens can be used for important public health and evolution genomic analyses, especially outbreak analysis and transmission reconstruction. Identifying this variation is typically achieved by aligning sequence reads to a reference genome, but this approach is susceptible to reference biases and requires careful filtering of called genotypes. Additionally, while the volume of bacterial genomes continues to grow, tools which can accurately and quickly call genetic variation between sequences have not kept pace. There is a need for tools which can process this large volume of data, providing rapid results, but remain simple so they can be used without highly trained bioinformaticians, expensive data analysis, and long term storage and processing of large files.

Here we describe Split K-mer Analysis (SKA2), a method which supports both reference-free and reference-based mapping to quickly and accurately genotype populations of bacteria using sequencing reads or genome assemblies. SKA2 is highly accurate for closely related samples, and in outbreak simulations we show superior variant recall compared to reference-based methods, with no false positives. We also show that within bacterial strains, where it is possible to construct a clonal frame, SKA2 can also accurately map variants to a reference, and be used with recombination detection methods to rapidly reconstruct vertical evolutionary history. SKA2 is many times faster than comparable methods and can be used to add new genomes to an existing call set, allowing sequential use without the need to reanalyse entire collections. Given its robust implementation, inherent absence of reference bias and high accuracy, SKA2 has the potential to become the tool of choice for genotyping bacteria and can help expand the uses of genome data in evolutionary and epidemiological analyses. SKA2 is implemented in Rust and is freely available at https://github.com/bacpop/ska.rust.

## Introduction

Pathogen genomes accumulate sequence variation over time, and mapping these differences to their genomes has proven invaluable for tracking outbreaks and informing public health interventions (Harris et al. 2010; Gardy et al. 2011; Grad et al. 2012; Quick et al. 2016). Genetic distances are a powerful additional source of information for inferring transmission events in outbreaks, allowing transmission events to be comprehensively ruled out where epidemiological links may suggest otherwise (Cori et al. 2018; Wymant et al. 2018). In particular, single nucleotide polymorphisms (SNPs) make up a substantial portion of the molecular events which generate genetic diversity among pathogenic strains. SNPs are ideal for inferring evolutionary relationships due to their high resolution, and the availability of tractable molecular models for phylogenetics.

From a practical perspective, identification of SNPs still relies on complex and computationally intensive pipelines usually based on read alignment, which requires significant bioinformatics expertise for tuning and maintenance, and large computing resources. Making reliable SNP calling quicker and easier would lower the barriers to routine implementation for transmission tracing, and assist academic scientists wishing to quickly investigate populations of pathogens before launching more involved analyses. Especially as volumes of pathogen genome data continue to grow (Hunt et al. 2024), tools which can make use of large populations without requiring excessive disk space or compute time can make the iterative cycle of investigation much more interactive and accessible. This is particularly the case for low-resource settings, which often have a high burden of bacterial diseases. Genome data is large and complex, even more so when dealing with populations. Unlike tabular data, which is easy to manipulate and plot, genome data has remained relatively undemocratic; its use is still mostly restricted to experts with good computational resources and training.

Identifying SNPs within a population traditionally involves aligning each sample’s sequencing reads against a reference genome. This involves multiple algorithms and pieces of software, so the resulting pipelines are often complex and sensitive to parameters, and may not necessarily be well-tuned to every pathogen, often restricting their usage to users with strong bioinformatics training and resources. Additionally, the choice of reference genome, read-alignment pipeline and its parameters can have a significant impact on SNP identification and influence downstream analyses (Pightling et al. 2014; Usongo et al. 2018; Bush et al. 2020; Walter et al. 2020). Differences in genomic composition and organisation between the input sequences and the chosen reference can generate misalignments leading to the identification of spurious SNPs (Landan and Graur 2009; Farrer et al. 2013; Hurgobin and Edwards 2017). Without sufficient coverage, this approach is more likely to call the reference variant, a phenomenon known as ‘soft reference bias’ (Paten et al. 2017). Alignment pipelines mitigate these errors using various alignment and SNP filtering criteria (e.g. matching alignment strands), excluding certain genomic regions or selecting a reference genome closely related to the strains of interest (Colquhoun et al. 2021). With typical mapping approaches, SNPs are totally missed when they occur in genomic regions that are either absent from or too dissimilar to the reference genome, a phenomenon known as ‘hard reference bias’.

An alternative approach, and the subject of our work, is to use subsequence probes i.e. k-mers to probe variation between samples, which avoids mapping and variant calling steps entirely. Odd-length k-mers can be further divided into two smaller fragments surrounding a variable middle base, forming a structure called ‘split k-mer’ (Gardner and Hall 2013; Harris 2018). In this structure, the left and right parts of the k-mer serve as local reference points to map the position of the middle base. If the same combination of left-right k-mers is found in another strain with a different middle base, homology between the two middle bases can be hypothesised and their difference interpreted as a SNP. The data conversion into split k-mers essentially creates independent local references for each genomic position of the strains of interest on the fly, enabling alignment-free and reference-free comparisons between them. Additionally, the split k-mers can be mapped to a reference sequence to impose a useful and interpretable ordering and coordinate system on the called variants.

Alignment-free algorithms can entirely avoid both forms of reference bias, so are well suited to bacterial species where these biases have a major effect on accuracy. The split k-mer approach does not use alignment during construction, so is robust to diversity from a reference – either avoiding references entirely as in ska align, or using the reference only as a coordinate system as in ska map. It is however important to note that cases of high diversity *among* samples cannot be addressed by SKA.

The split k-mer approach was first introduced through the kSNP program (Gardner and Hall 2013; Gardner et al. 2015; Hall and Nisbet 2023). We previously implemented an enhanced version of this approach in the SKA (’split k-mer analysis’) software, hereafter referred to as ‘SKA1’. SKA1 offered improvements in efficiency and flexibility and the ability to map detected SNPs to a reference genome (Harris 2018). SKA1 has since been successfully employed in genomic studies of a large range of bacterial pathogens (Becker et al. 2021; Gladstone et al. 2021; Morris et al. 2022; Chew et al. 2023), and has independently been shown to be superior to limited resolution gene-by-gene approaches for mapping transmission (Maechler et al. 2023). There remains a need for these variant calling methods which scale to the size of modern datasets, and which produce high quality results without needing manual testing and adjustment, and avoid soft and hard reference bias common in analysis of bacterial populations (Bush et al. 2020; Colquhoun et al. 2021; Valiente-Mullor et al. 2021).

We have therefore re-engineered and improved our implementation to develop SKA2, which we designed to be faster, more flexible, and easier to maintain. SKA2 is intentionally designed as a tool which is quick and easy to use. Academics can use SKA2 to quickly analyse data and test hypotheses. In public health and clinical settings, SKA2 can rapidly determine whether an isolate is from an outbreak quickly and without the need for dedicated high-performance computing. At the same time, users have access to a method which is as accurate as the more complex, detailed analyses for the particular question they have.

## Material and Methods

### Split k-mer encoding

A split k-mer is broadly defined as a sequence of length k, containing one or more contiguous unspecified (‘wildcard’) bases which can match with any character (Harris 2018). Throughout this paper we use a more specific definition, which is an odd length k-mer where the middle base is allowed to vary (i.e. the only wildcard position is the middle one). Other choices, including having the wildcard position as the final or first base would be possible, but including the base within the k-mer ensures only SNPs can be tagged, as a variable base at the end may be the start of a larger genetic event. We therefore opt for the middle base following the precedent of SKA1. For example, with k = 11 the split k–mer would be XXXXX-XXXXX, where X={A,C,G,T} and ‘-’ is unspecified. The methods used in SKA2 are demonstrated with an example with k=11 in Figure 1.

**Figure 1:**
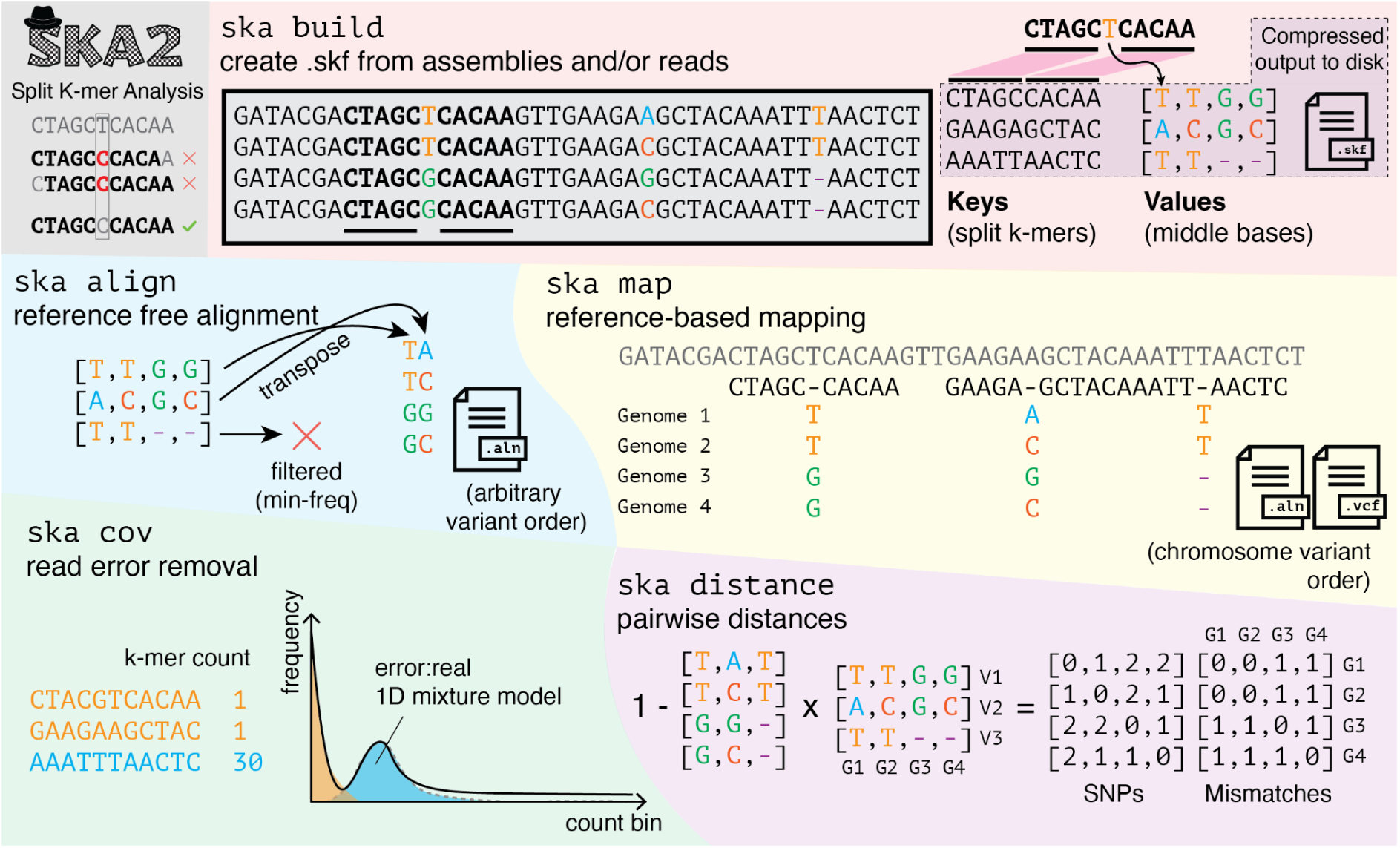
Overview of functions and methods in SKA2. Split k-mers allow matching variant positions, whereas contiguous k-mers mismatch any variation. ska build creates split k-mer dictionaries from input sequence data. The example shows four sequences which are aligned and on the same strand for clarity, but in real input data neither is necessary. Split k-mers are used as keys and their middle bases are stored in lists. This dictionary is compressed using snappy to make .skf files. ska align makes reference-free alignments with no coordinate system by writing out the middle bases, applying filters on frequency of missing data, constant sites and ambiguous sites. ska map makes reference-based mappings as .aln or .vcf, with the same coordinate system as the reference. In both modes the conserved sites are also written out, but not shown for clearer visualisation. ska cov counts k-mers and fits a mixture model to find a threshold for count when using reads as input to ska build. ska distance calculates SNP distances and mismatches between samples by multiplying the middle base matrix by its transpose. The cluster_dists.py script can be run on this distance matrix to make a phylogeny, single linkage clusters with a provided threshold, and a Microreact visualisation. Operations to merge, delete samples and split k-mers, and write out the contents of .skf files are also implemented, but not shown.

To count the split k-mers in an input fasta file we read each new base in turn and use the following commonly-used mapping to two bits: {A, C, G, T} = {00, 01, 11, 10} (Drezen et al. 2014). This can be efficiently converted into the upper and lower half of the split k-mer using bitshift and masking operations, rolling through the input string rather than rebuilding the split k-mer at every position. To match the machine word size, for k <= 31 the known bases are packed into 64-bits, for k <= 63 the known bases are packed into 128-bits. The default is to consider both of the integer representations of the k-mer and its reverse complement and keep only the smallest, which is consistent irrespective of which stand the k-mer was actually counted on. If the strand is made to be consistent between samples ahead of time (e.g. single-stranded viruses or reference sequences), this step can be skipped.

We store the middle base in a single byte, and thereby support IUPAC uncertainty codes. Any observation of the same split k-mer contributes equally to the uncertainty in the middle base (i.e. a single duplicate observation adds that base as a possibility), and the specific counts of different middle bases are ignored. Due to the split k-mer necessarily containing an even number of fixed bases, it is possible for the fixed part of a split k-mer to be its own reverse complement (a palindrome). In this case the strand of the middle base is ambiguous, so we keep both the observed base and its complement as observations using uncertainty codes.

The fundamental data structure we use in SKA2 for each input sequence is a hash map (dictionary), with the split k-mers as the keys, and their middle bases as the values for the entry. For example, using k = 11 and ignoring the reverse-complement sequences for simplicity, the sequence CTAGCTCACAAGT would have dictionary entries {CTAGCCACAA: T; TAGCTACAAG: C, AGCTCCAAGT: A}.

### Creating split k-mers from short read sequencing data

When using fastq files as input, ska assumes these have been generated from error-prone short read sequencing. We implement two filtering options to prevent sequencing errors from entering the split k-mer dictionary. The first is a minimum quality score for reported read quality (default = 20), which can be applied at the middle-base only (’middle’) or across the entire split k-mer (’strict’, the default).

The second is a minimum observation count (default = 5). Keeping track of k-mer counts, even in relatively small sequencing experiments, takes >90% of the runtime, so filtering on the minimum observation count is an important optimisation target. For simulated 150bp paired-end reads from *Mycobacterium tuberculosis* at 60x coverage, a baseline of keeping counts in a dictionary takes 99s and uses 2.4Gb of memory. We evaluated numerous alternative approaches, and found that a two-stage approach, using a blocked-Bloom filter to remove singletons and then a dictionary to count k-mers appearing at least twice was the fastest, least memory-consuming, and most accurate insofar as it gave no false positives or false negatives with the default count threshold. This is similar to the approach taken by bifrost (Holley and Melsted 2020). The hashes in the Bloom filter are calculated using ntHash (Mohamadi et al. 2016), the filter uses a word-aligned block with 12 bits per block (∼1% FPR) and is 2^27 blocks wide (∼200M bits) (Qiao et al. 2014). For the same simulated sample, this approach takes 46s and uses 1.6Gb of memory.

To select a suitable minimum k-mer count we implemented a coverage model (ska cov) which fits a mixture model to the count frequency distribution of contiguous k-mers. There are two advantages of a model-based approach over a simpler method such as choosing the minimum count bin: the model is guaranteed to have a single global minimum, and can be fit to find a single threshold even with noisy data.

We used an observation model which is a mixture of error k-mers and real k-mers. The error component probability density is a Poisson count with mean one, the real k-mer probability density is a Poisson count with mean of the coverage *c*. Their mixture with *ω* as the proportion of error k-mers has likelihood:

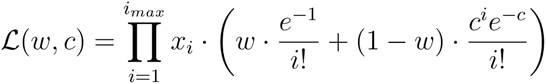

Where the data *x_i_* is the number of split k-mers counted *i* times. We fit *ω* and *c* by maximising the likelihood. We do this using the BFGS algorithm (Fletcher 1970) and deriving analytic expressions for the gradient 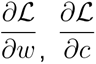. The count cutoff is selected by picking the first integer value of *x_i_* at which the responsibility of the non-error component is greater than the error component. An example of observed coverage counts and the k-mer model fit is shown in supplementary figure 1.

### Split k-mer genotyping

To create an alignment, the multiple dictionaries from each sample are merged into a single dictionary. The values in this dictionary are then lists of the middle bases for each sample, using a gap character ‘-’ if the split k-mer was not observed in that sample. We implement two operations on this merged dictionary: append (to add a single sample) and merge (to combine two merged dictionaries). Using these operations with the same algorithm as a parallel merge sort it is possible to support parallel construction and merging. As the merge operation is slower than append this only gives a speedup when construction of the initial dictionaries is slow, i.e. with read data. With two threads on a dataset of 28 samples with 3.2M split k-mers our approach yields a speedup of 1.7x. The ska build command combines this merge with the encoding for multiple samples as described above. By doing this we minimise the memory and disk requirements and simplify the command line use. It is also possible to parallelise the build over samples using a program such as GNU parallel, followed by a single merge. This process gives close to a 100% efficient speedup, at the expense of increased memory usage and more temporary disk space.

In the default, reference-free mode, ska align, each aligned column is a list of the middle bases in this merged array. Therefore creating the output alignment simply consists of writing the transpose of the dictionary values. Options to filter the sites are available: a minimum number of observations (i.e. non-missing middle bases); remove any ambiguous sites; replace any ambiguity with ‘N’; no constant sites; allow ‘-’ in otherwise constant sites (useful for low coverage samples). These columns are in an arbitrary order, as in the hash table, and do not represent a physical position in the chromosome.

In the reference-based mode ska map, the algorithm iterates over the split-k-mers in the input reference sequence, searching for each one in the merged dictionary. If a match is found, the middle base at that position is written, otherwise a missing base ‘-’ is output. Extra logic is needed to write the matching parts of the split k-mers, keep track of multiple chromosomes, and mask mapped but repeated split k-mers with N (if requested by the user). We have implemented rigorous unit-testing to ensure this process works correctly. Output as either FASTA or VCF is supported.

### Split k-mer files and their supported operations

We save split k-mer files (.skf) to disk for reuse, addition, or alternative filtering. These consist of the list of split k-mer keys, an array of the middle bases, the count of samples in which each split-k-mer has occurred, the sample names, and some metadata (k-mer size, version, and number of bits used for k-mer keys). To generate these we serialise the split k-mer dictionaries and compress them using snappy, which adds minimal time to read/write operations and achieves a compression ratio of around 10x. Using .skf files we support merging with another .skf file (ska merge), deleting named samples (ska delete), and deleting specified split k-mers or refiltering the split k-mers (with ska weed). The number of k-mers command (ska nk) can be used to output the contents of .skf files to the terminal, including the total number of split k-mers and the split k-mer observed in each sample, which can be a useful quality control metric (e.g. elevated in the case of contamination). The same command can optionally output, in text form, the split k-mer sequences and the middle bases of all samples, which can be used for processing in other bioinformatic tools.

### Sample distances and clustering

We implemented a ska distance command which iterates over every pair of samples to calculate the number of SNPs different between the samples. For each matching split k-mer, the probability vectors of their middle bases are multiplied to get their overlap. For example, if a sample has either a C or a G as a middle base, we give a 50% weight to each one, and then compute the probability of a match by multiplying the probabilities. Then, 1 - the probability of a match gives the probability of a mismatch (which is 1-distance). These probability vectors are set by the IUPAC codes, with the probability equally divided among possible middle bases. For assemblies and small numbers of repeats this is reasonable, but will be less effective than a full variant caller which tracks the count of each observation. The number of missing sites is also reported as mismatches. To fully replicate and extend the functionality of SKA1 we also wrote a downstream python script which clusters samples using single linkage at a given SNP threshold, makes a neighbour-joining tree with rapidnj (Simonsen et al. 2008), and creates a visualisation of the clusters, tree and single linkage graph. This is displayed interactively using Microreact (Argimón et al. 2016), an example is shown in supplementary figure 2.

### Implementation and comparison to SKA1

SKA2 is implemented in rust and is freely available at https://github.com/bacpop/ska.rust (Apache 2.0 license) with installation also possible via crates.io, bioconda or directly from source. Documentation, including API code for reuse in other software is available at https://docs.rs/ska/latest/ska/.

A tutorial is available at https://www.bacpop.org/guides/building_trees_with_ska/. We designed SKA2 so that the majority of SKA1 functions would also be available, but a full list of differences is given in the software documentation. The code runs automated tests and ensures at least 90% test coverage of the code. Each version is automatically compiled and so can be installed as a static binary, using the rust toolchain to compile locally, or via bioconda. All tests in this paper used version 0.3.5.

Some features of SKA1 were not implemented in SKA2. The maximum k-mer size is 63 (to fit into a 128-bit integer, the maximum native size supported by Rust) rather than unlimited – our testing showed that longer k-mer sizes did not yield improvements (Figure 2). The annotate function was not implemented, as its functionality is covered by bedtools (Quinlan and Hall 2010). The unique function can be parsed from the output of the dictionary, and was not made into a separate function. The type function has been superseded by k-mer tools such as StringMLST (Gupta et al. 2017) and BioHansel (Labbé et al. 2021).

**Figure 2:**
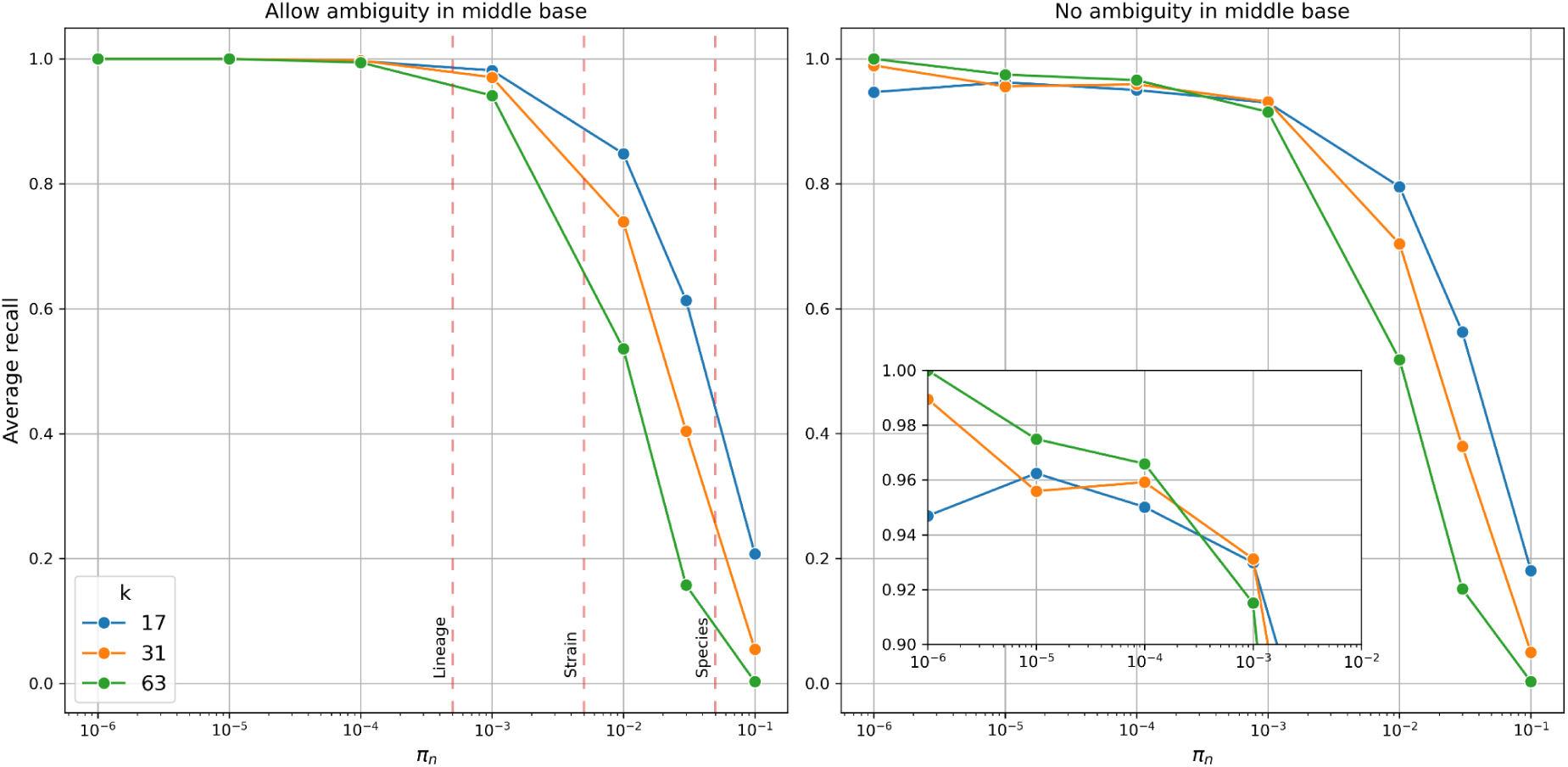
Average recall of SKA2 in simulations across increasing sequence divergence between a pair of sequences (π_n_ or SNPs per site). Lines show recall using different split k-mer lengths *k*. Left: recall when allowing ambiguous bases, showing typical divergence thresholds used to define species, strain and lineage boundaries. Right: recall when requiring exact matches of the middle base, with inset showing recall over the within-lineage range.

SKA1 has the option, through ska merge, to invert the keys and values in the merged dictionary. In this format, each unique pattern of middle bases appears only once in the keys, and the corresponding split k-mers in a vector of the values. For closely related samples this is effective at reducing redundancy; the above example has only 1163 unique patterns from 3264141 split k-mers, so the output compresses to 34Mb. We did not implement this in SKA2 as the gain over the current compression approach is marginal and increases processing time.

### Split k-mer simulations

We simulated a mutated sequence along a single branch using the Gillespie algorithm (Gillespie 1977), following a similar approach to that taken in phyloSim (Sipos et al. 2011). We used a GTR+Gamma+Invar rate model plus short indels, with parameters previously estimated for *Streptococcus pneumoniae (Lees et al. 2018)*, using reference ATCC 7000669 (Spn23F) as the starting sequence (Croucher, Vernikos, et al. 2011). We tracked the exact expected split k-mers for each substitution in the simulation and calculated power as the proportion of these detected by running ska align between the starting sequence and the mutated sequence. We calculated power requiring an exact match of the split k-mer and the correct middle base, and also a more relaxed mode that just checks the flanking sequence, tolerating ambiguity at repeat sequences. Indels are ignored in the power calculation, but may reduce power if they interrupt a split k-mer. The code for this analysis can be found at https://github.com/bacpop/ska_simulations (v0.1.0).

### Outbreak simulations

We simulated outbreak clusters of *S. pneumoniae* and *M. tuberculosis* using Transphylo v1.4.10 (Didelot et al. 2017) with simulation parameters provided in Supplementary Materials. We then simulated sets of mutations along these phylogenies using phastSim (De Maio et al. 2022). We choose a mutation rate of 10 and 1 mutations per genome per year for *S. pneumoniae* and *M. tuberculosis* respectively, which correspond to upper estimates of the mutational rate of these two species (Chewapreecha et al. 2014; Menardo et al. 2019). For *S. pneumoniae*, we also simulated indels of 100 bp at a rate of 0.5 indel per year to mimic accessory genome changes as in (Lees et al. 2019). phastSim command lines are provided in Supplementary Materials. SNPs and indels were then randomly inserted in genome assemblies using a custom python script to recreate the genome of each sample composing the outbreaks. We finally simulated Illumina HiSeq 2500 paired-end sequencing reads of 150bp from the modified genome assemblies at a 60x coverage using ART (Huang et al. 2012).

We used the variant calling pipelines Snippy v4.6.0 (https://github.com/tseemann/snippy) and BactSNP v1.1.0 (Yoshimura et al. 2019), both with default parameters. The command lines of the ‘BWA+BCFtools’ read-alignment pipeline are provided in Supplementary Materials. Briefly, reads were aligned using BWA-mem v0.7.17 (Li 2013), SNPs were called using BCFtools v1.10.2 (Li 2011) and then filtered using the following criteria: a minimum of 10x coverage, supported by at least one read forward and one read reverse, and a minimum allele frequency of 0.75. SNP alignments obtained from all read-alignment pipelines were finally trimmed to only retain positions with a maximum of 10% missing data, considering degenerate nucleotides as missing. Runtime and memory consumption were measured on an Intel Xeon Platinum 8360Y CPU with 10GB RAM memory. The parameters and commands used in the simulations are listed in supplementary data 1.

False positive and false negative SNPs were identified by comparing the positions of the inferred SNP alignments to those of the expected SNP alignment using a custom Python script available and described at https://github.com/rderelle/compareALI. Phylogenetic analyses were performed using IQtree2 v2.2.2.7 (Minh et al. 2020) under the GTR+ASC model and near-zero branches collapsed into polytomies (‘--polytomy’ option). Distances between phylogenetic trees were computed using the R package TreeDist v2.7.0 (Smith 2023).

### Recombination analysis

We used ska build, with both k = 17 and k = 63, followed by ska map with options to mask repeats and any ambiguous bases to create an alignment of 240 samples against the Spn23F reference (Croucher et al. 2009). We used gubbins v3.3.1 (Croucher et al. 2015) turning off gap filtering (--filter-percentage 100.0) with otherwise default parameters (no outgroup, RAxML as tree builder, five iterations). We compared the density of recombinations across the genome to the originally published analysis using phandango (Hadfield et al. 2017) which includes this data as a default example set.

### Genome data

28 *Listeria monocytogenes* samples from a study of bacterial meningitis, taken from the largest PopPUNK cluster, were used for testing and comparison with SKA1 (Kremer et al. 2017; Lees et al. 2018; Lees et al. 2019). The 288 *E. coli* genome assemblies used for the online analyses were extracted from the genometrackr database (Allard et al. 2016), and all correspond to a single strain in the PopPUNK database (Lees et al. 2019). Their accession numbers are provided in supplementary table 2. For the gubbins analyses, 240 *S. pneumoniae* genomes from the PMEN1 strain were used (Croucher, Harris, et al. 2011).

## Results

SKA2 can be used to genotype SNPs in sets of populations using ska align. These variants are unordered so are only suitable for constructing phylogenies or transmission detection. The ska distance tool can also be used to create possible transmission clusters by applying a SNP cutoff. In outbreak settings, this mode is highly accurate and can be used rapidly with either or both read (60x speedup) and assembly data (190x speedup) as input. Despite the simple approach to SNP genotyping, in its primary use case of outbreak settings it outperforms traditional read mapping approaches that dominate this type of analysis. In the first section we show that this holds within related bacterial samples (corresponding to a typical strain or sequence-type cluster definition), then in the second section use simulated outbreak data to compare to read mapping approaches. This approach also avoids the need for selection of an appropriate reference, which requires expertise, and introduces hard- and soft-reference bias which lower the sensitivity of variant calling.

The mapping approach of ska map gives mapped variants a chromosome coordinate. Although in small sample sets of possible transmission this has less power than ska align, mapping has advantages where the order and reference-based functional interpretation of variants needs to be kept, which we demonstrate in an example for recombination detection in the third section below. For large sample sets where diversity may be larger, ska map remains scalable and robust.

Tools to investigate shared k-mer content ska nk on a per-sample basis are useful quality control, and can rapidly determine the diversity of a population. Using ska weed to remove or keep chosen sequences can also rapidly be used to detect the presence of genetic elements such as transposons. In the final section, we show these methods can be used to create scalable split k-mer files for mapping with large and continuously growing sample collections.

Compared with SKA1, SKA2 showed improved runtimes and smaller file sizes, a major objective of our reimplementation. For a 3Mb bacterial assembly (*Listeria monocytogenes*) the build process of SKA2 takes 0.1s on average, producing a .skf file 16Mb in size. In SKA1, the process takes 2.8s and produces a .skf file 30Mb in size. For 28 samples of *L. monocytogenes* from the same strain, the .skf file is 38Mb and takes 5s to create. In SKA1, these files are 870Mb and take 194s to create and merge (∼40x speedup). For a set of simulated reads, SKA2 build takes 46s and uses 1.4Gb of memory. In SKA1 this takes 120s (∼2.5x speedup) and 0.6Gb memory. A major advantage of SKA2 over SKA1 is therefore the optimised and streamlined combined build and merge step, and the resulting smaller files. This increase in speed is due to a faster dictionary implementation, optimised file parsing, and use of a faster serialisation process. Parallelisation is not directly supported in SKA1.

### Split k-mers accurately and quickly genotype closely related samples

Before using SKA2 to analyse bacterial populations, we first used two-sample simulations to evaluate the power and limitations of the split k-mer genotyping approach. We first note that more complex variation, such as insertions, deletions (indels), rearrangements or copy number variation is ignored by SKA2 entirely. However, as typically only SNP calls are used in reconstructing evolutionary history (Lees et al. 2018) due to their mostly vertical inheritance, and existence of corresponding effective models of molecular evolution, we focus on these variants entirely.

As SKA2 is alignment free, it is unaffected by soft reference bias (more likely to call the reference base) (Colquhoun et al. 2021). ska align is also robust to changes in sequence content, so unaffected by hard reference bias (missing accessory genes). However, split k-mers can still miss SNP variant calls, causing false negatives, for the following reasons:

- When using assemblies as input, variants closer than half the split k-mer length to a chromosome or contig boundary cause the flanking bases to be too short to be enumerated in the dictionary. When using reads as input a similar problem occurs at the ends, in the case of very low coverage data.
- Two or more SNPs within the half k-mer arm length (i.e. (k-1)/2) of one another cause a mismatch in the flanking bases.
- Similarly, indels or rearrangements can interrupt split k-mers.
- Repeated split k-mers cannot be uniquely mapped, so their middle bases contain ambiguity from all observations of the split k-mer.

We also note that doubly (or more) mutated sites, occurring more frequently at larger distances, only count the most recent variant. In simulations it is possible to detect this as an error from the reference sequence, but is not an issue with variant calling methods themselves.

Unremoved sequencing errors in reads (due to poor filtering) or assembly errors may introduce false positive variant calls prior to input to the SKA2 data structure, but split k-mers themselves introduce effectively zero further false positive variant calls. We therefore focused on false negatives here. False positives are assessed further in the outbreak simulations below.

We used simulations to evaluate the relative importance of these effects across evolutionary distances (Figure 2, left panel). We found that the major effect on calling accuracy was variants occurring closer than the k-mer distance. Indel rate had a negligible effect on power, as it was no more likely to interrupt a split k-mer than a nearby SNP (supplementary figure 3). SKA is intended to be used for closely related samples, and shows good power in this range. Within a lineage (divergence ≲ 0.0005 SNPs per site, in some species referred to as a clone) the recall is >99%. Outbreaks are often much more closely related than this as many species acquire only a few SNPs per year (Duchêne et al. 2016), showing that SKA2 is a reliable genotyping tool for this purpose.

Within a bacterial strain (divergence ≲ 0.005 SNPs per site) this drops slightly to around 90%, and across an entire bacterial species (divergence ≳ 0.05 SNPs per site), the recall drops further, to between 10-50%. When ignoring repeats, defined here as any split k-mer with multiple observations, shorter k-mers had consistently higher power as they are proportionally less susceptible to multiple close SNPs. We therefore recommend that SKA2 is used only within bacterial strains and for this purpose using a short k-mer length.

The main effect on accuracy is the resolution of repeats (Figure 2, right panel), though we note that this is a strict criterion, as most phylogenetic software allows uncertainty in alignments. Requiring exactly resolved repeats results in an average recall to around 95%, though using longer k-mers improves this up to around 98%. Split k-mers show the archetypal sensitivity/specificity tradeoff for length: at small divergences longer split k-mers are preferred as they are more specific, due to uniquely mapping more sequences by spanning shorter repeats; at larger divergences smaller split k-mers are preferred as they are more sensitive, due to mapping more nearby SNPs.

### SKA2 outperforms read-mapping approaches in simulated outbreak analyses

Having confirmed that split k-mers are most effective for closely related examples, we next evaluated the performance of SKA2 in simulated outbreaks, comparing SKA2 to traditional read-alignment methods. We analysed simulated outbreaks of *S. pneumoniae* (12 samples; 87 SNPs) and *M. tuberculosis* (30 samples; 38 SNPs). Each simulated outbreak was replicated five times, leading to 20 simulated outbreaks per species. We generated sequences at the tips of each expected phylogeny from the transmission series, then created simulated short-read sequences from this truth set of full length sequences. We tested the effect of reference bias by using references of increasing divergence from the root genome (ATCC_700669 and H37Rv for *S. pneumoniae* and *M. tuberculosis* respectively). We compared SKA2 to the popular read-alignment tool Snippy, the hybrid assembly-mapping pipeline BactSNP and a custom-made SNP-calling pipeline referred hereafter as BWA+BCFtools. We used SKA2 in both reference-free mode (align) and its reference-based mode (map).

For *S. pneumoniae* outbreaks, read-alignment pipelines showed increasing numbers of false negative SNPs when more distant outbreaks to the reference genome were analysed, highlighting the impact of differential genome composition between the analysed outbreaks and the reference genome (Figure 3). In contrast, ska align, showed relatively stable numbers of missed SNPs, regardless of the origin of the outbreak: SKA2 missed similar numbers of SNPs to read-alignment pipelines when the analysed outbreak corresponds to the reference genome (here from the strain ATCC 700669), but significantly less when outbreaks isolates were much less similar to the reference genome. None of the SNP inference methods produced false positive SNP when outbreaks derived from the ATCC 700669 reference genome were analysed. However, Snippy, BWA+BCFtools and BactSNP produced on average 89.1, 6.4 and 0.2 false positive SNPs respectively when more distant outbreaks were analysed. The low specificity of Snippy has already been characterised in benchmark studies (Yoshimura et al. 2019; Falconer et al. 2022). In contrast, SKA2 did not produce any false positive SNP in any of these analyses. We then compared the phylogenetic trees obtained from all these SNP alignments to the tree obtained from the set of simulated SNPs. With the exception of outbreaks derived from the ATCC 700669 genome, the phylogenetic trees obtained from BactSNP and SKA2 SNPs were found to be more similar to the expected tree than those derived from SNPs inferred by Snippy and BWA+BCFtools read-alignment pipeline.

**Figure 3:**
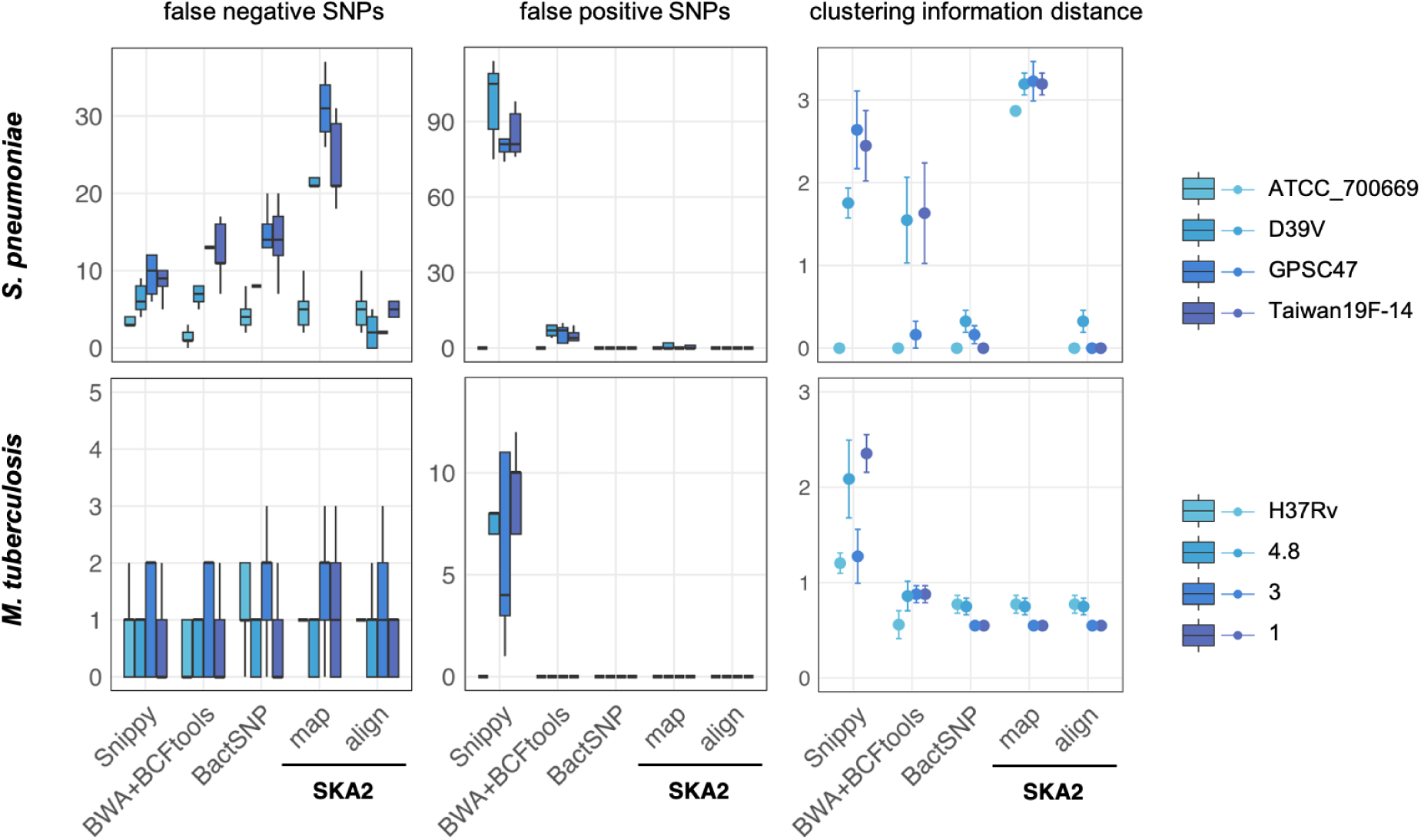
Results obtained from the analyses of simulated outbreaks showing recall (false negatives), false positives and clustering information distance from the four different tools. “map” and “align” refer to the SKA2 functions used to generate SNP alignments. References of increasing distance (darker blue) from the source of the outbreak were used to evaluate reference bias. The error bars in the CI distance plots correspond to the 95% confidence interval calculated from 10 values (two phylogenies were obtained from each SNP alignment using two independent maximum-likelihood runs). The numbers 4.8, 3 and 1 in the legend correspond to the names of *M. tuberculosis* lineages.

For *M. tuberculosis*, which is characterised by a highly conserved genome across lineages (Comas et al. 2013; Gagneux 2018), SKA2 still demonstrated similar or superior performance when compared to read-alignment methods, as illustrated in Figure 3. The numbers of false negative SNPs were found to be similar between all methods, and BactSNP and SKA2 were the only methods to not produce any false positive SNP (on average 7.6 and 0.1 false positive SNPs generated by Snippy and BWA+BCFtools in non-H27Rv outbreaks). The phylogenetic trees obtained from BWA+BCFtools, BactSNP and SKA2 SNPs were all found to be similarly distant to the expected tree while those obtained from SNPs inferred by Snippy were found to be more distant.

Using ska map gives very low, but not always zero, false positives. We investigated these false positives and found them to be present in tandem repeats. In cases where the flanks of a split k-mer can be mapped in multiple overlapping places, it is ambiguous whether the mapped base should be overwritten by the conserved flank or the middle base (for consistency of reporting, the code always chooses the latter option). Filtering on repeat regions removes these false positives, but increases false negatives. For a purely phylogenetic or transmission analysis, the lower recall (which drops further with more distant references) is less desirable. However, when using many samples the ability to map them one at a time can still be useful – with ska align the entire intersection of split k-mers must always be kept, which leads to large memory use with diverse samples to keep low frequency (and typically unused) split k-mers. In such sample sets, using the default of 80% presence can also lead to loss of k-mers in ska align which can be usefully retained by ska map.

We repeated the SKA2 analyses to assess the impact of split k-mer size on its SNP detection. Using k-mer sizes ranging from 21 to 61 nucleotides, in increments of 10, we observed similar numbers of identified SNPs between k=21 and 51 for *S. pneumoniae*, followed by a sharp decrease at k=61 (supplementary figure 4). For *M. tuberculosis*, we observed a slight increase in the number of identified SNPs from k=21 and 31 followed by a plateau and again a sharp decrease at k=61 (supplementary figure 4). Then, we evaluated the performance of SKA2 in low sequence coverage settings, from 60x coverage used in the previous set of analyses down to 10x coverage, in decrements of 10. We observed a sharp decrease in the sensitivity of SKA2 at 20x coverage when filtering settings are not adjusted (average sensitivity of 0.93 to 0.43, and 0.96 to 0.31 for *S. pneumoniae* and *M. tuberculosis* outbreaks respectively; supplementary figure 5). Reducing the minimum count threshold for the detection of split k-mers from five to three restored the sensitivity of SKA2 to 0.89 and 0.93 at 20x coverage for *S. pneumoniae* and *M. tuberculosis* outbreaks respectively, which correspond to sensitivity levels obtained by Snippy at the same coverage. These analyses also revealed a small rate of false positive SNPs inferred by SKA2 at 30x and lower coverages due to unfiltered sequencing errors, though on average less than 0.5 false positive SNP per simulated outbreak (supplementary figure 5). These spurious SNPs might be caused by differential distribution among samples of split-k-mers corresponding to duplicated genomic regions due to stochastic variations of coverage.

We also recorded runtimes and maximum memory consumption of SKA2, SKA1 and read-alignment pipelines in these outbreak analyses. SKA2 was found to be by far the fastest method: two to four-fold faster than SKA1, 14 to 20-fold faster than the read-alignment pipelines Snippy and BWA+BCFtools, and 60-fold faster than the hybrid pipeline BactSNP (Table 1). Despite the necessity to store all split k-mers in memory, SKA2 also displayed modest maximum memory consumption values, which were similar to those observed with SKA1 and other variant calling pipelines. Finally, the skf files produced by SKA2 were on average only 24 and 53 Mb in size per outbreak for *S. pneumoniae* and *M. tuberculosis* respectively (31 and 66 Mb for SKA1 output files), while the BAM files produced by the read-alignment pipelines were on average 91 and 185 Mb in size per sample for *S. pneumoniae* and *M. tuberculosis* respectively. SKA2 disk usage was therefore found to be 40-100 times lower than traditional read-alignment.

**Table 1:** Runtimes obtained on one CPU (in minutes) and maximum memory consumptions (in Mb) in the analyses of simulated outbreaks. Values represent averages over 20 replicates. ”n” represents the number of samples per outbreak. The lowest values are highlighted in bold.

|  | *S. pneumoniae*<br>(n=12) |  | *M. tuberculosis*<br>(n=30) |  |
| --- | --- | --- | --- | --- |
|  | Runtime<br>(minutes) | Memory<br>(Mb) | Runtime<br>(minutes) | Memory<br>(Mb) |
| BactSNP | 59.6 | 1955 | 330.1 | 2059 |
| Snippy | 20.1 | 338 | 90.9 | 1434 |
| BWA+BCFtools | 19.8 | **323** | 77.4 | **656** |
| SKA1 align | 2.3 | 424 | 20.1 | 865 |
| SKA2 align | **1** | 443 | **5.5** | 893 |

We finally recorded runtimes and maximum memory usages of SKA2 using a real-world dataset composed of fastq files from 100 isolates of *S. pneumoniae* belonging to the IC1 cluster (Croucher et al. 2014). By repeating the analyses (ska build at k=31 and ska align functions) in increments of 10 isolates, we found that SKA2 runtimes increased nearly linearly with the number of isolates (Figure 4). In contrast, and as expected, its maximum memory usage was found to be dependent on the total number of split k-mers extracted from the sets of isolates.

**Figure 4:**
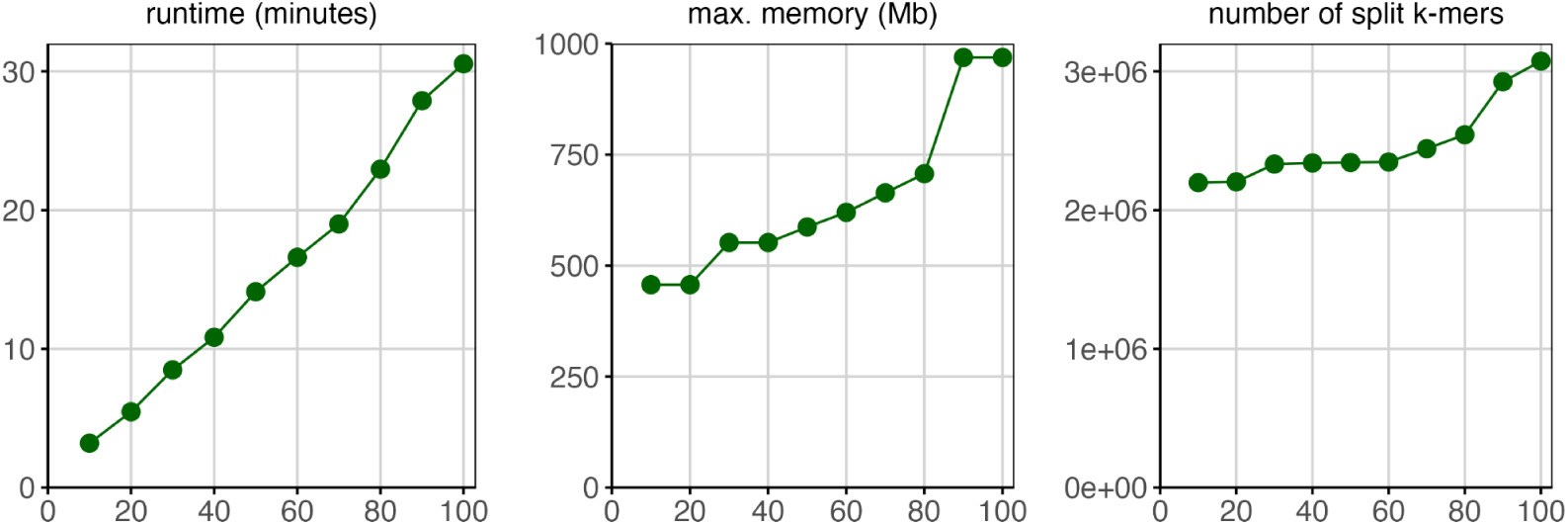
Empirical scaling of SKA2 computational efficiency using increasing block sizes from 100 isolates of *S. pneumoniae* IC1 cluster. The numbers of split k-mers represent the total numbers of split k-mers contained across all samples.

### SKA2’s map can be used effectively to detect recombination with assemblies as input

We show above that for phylogenetics, using ska align is superior to the reference-based approach of ska map. However the all-versus-one approach of ska map offers several key advantages: a unique comparison point based on which all variants are characterised, some functional interpretation by using the annotation of the reference, a well-defined coordinate system, computational scalability which can be trivially parallelised, and flexibility as new samples can be added to an already processed dataset.

For some purposes, ordering and spacing variants along the chromosome is necessary. This is typically for uses where the actual physical molecule structure is important to maintain local linkage disequilibrium. One example is recombination detection, which typically finds windows of locally elevated variant density which are more likely the result of transfer from a relative than local hypermutation (Croucher et al. 2015; Didelot and Wilson 2015; Mostowy et al. 2017). Another is genome-wide association studies, where local linkage disequilibrium is used to define distinct signals (Lees et al. 2016).

Anecdotally, a popular use of SKA1 has been to create ordered SNP alignments against a reference genome for use in recombination detection. This mode of input generation is also officially supported in the gubbins package (Croucher et al. 2015). Given that a limitation of split k-mers is that SNPs closer than the k-mer length will be missed, we tested that this is still sufficient to identify regions of elevated SNP density and produce correct results with recombination detection methods, Using 240 genome assemblies as input, SKA2 is able to rapidly create an input alignment against a reference chromosome (56s; 3.6Gb RAM), around ∼190x faster than using snippy on all samples (178 min). Using this as the input to gubbins gives similar recombination signals to the original results (supplementary figure 6), which were based on short read mapping (similar to the BWA+BCFtools pipeline). As in the original study, the major peaks spanning prophage MM1-2008φ, the mobile element ICESpn23FST81, and the *cps* operon were all detected. The only major locus not detected was the *psrP* gene. This gene mostly consists of short 2-3 amino acid serine-containing repeats, and hence this low complexity repeat sequence that is longer than the split k-mer length cannot be mapped by SKA2 (figure 2). This repeat is longer than 63 bases, so even using the maximum k=63 is unable to recover these SNPs.

### Split k-mers can be filtered to allow ‘online’ outbreak analysis with large numbers of samples

Noting the computational efficiency of SKA2, in particular with respect to file sizes, we then tested the suitability of SKA2 for ‘online’ (or serial) analysis where genomes are added to the existing in batches, rather than the entire dataset being reanalysed. This is the dominant mode by which genome data now arrives for bacterial pathogens, and supporting analysis in this manner will lower the computational burden, particularly relevant for democratising access to rapid outbreak analysis. We also consider these results in the context of making ‘reference’ .skf files available on a website (such as pathogen.watch) so users can use them to contextualise their new genomes. To be used for interactive internet tools, file sizes ≲10Mb are desirable. To efficiently store large amounts of data, we propose reducing the size of skf files by filtering split k-mers using the ska weed function. Here, we tested the power lost using an online approach by using iterative analyses of 288 *Escherichia coli* genome assemblies, in which the skf file generated from the new genomes is merged to the previous skf file, then filtered to remove constant or rare split k-mers. In absence of ground truth, we considered in these analyses the numbers of SNPs obtained from standard SKA2 analyses (i.e., without split-k-mer filtering) as reference.

Initially, we added *E. coli* genome assemblies in random batches of 10, employing a split k-mer filtering based on consistently missing split k-mers (i.e. removing any rare split k-mers; parameter min-freq set to 0.8). The number of SNPs found at each iteration closely matched the results obtained through standard SKA2 analyses (with a maximum difference of 13 SNPs out of 16,905 SNPs at 100 genomes). Simultaneously, the size of skf files was substantially reduced, ranging from 2 to 6.4-fold reductions after filtering in the first and last iterations, respectively (Figure 5), reaching a file size of 212 Mb for 288 genomes. For comparison, the 288 unprocessed and gzipped genome files represent a combined 454 Mb of data. Similar outcomes were observed when adding genomes in batches of 50, and starting with a dataset of 200 or 250 genomes followed by additions in batches of 5 or 1. These analyses affirm that SKA2 would be suitable for use in an online mode, and even suitable for use in a web browser tool, as it can proficiently store substantial genomic data with negligible information loss, facilitating the analysis of ongoing outbreaks.

**Figure 5:**
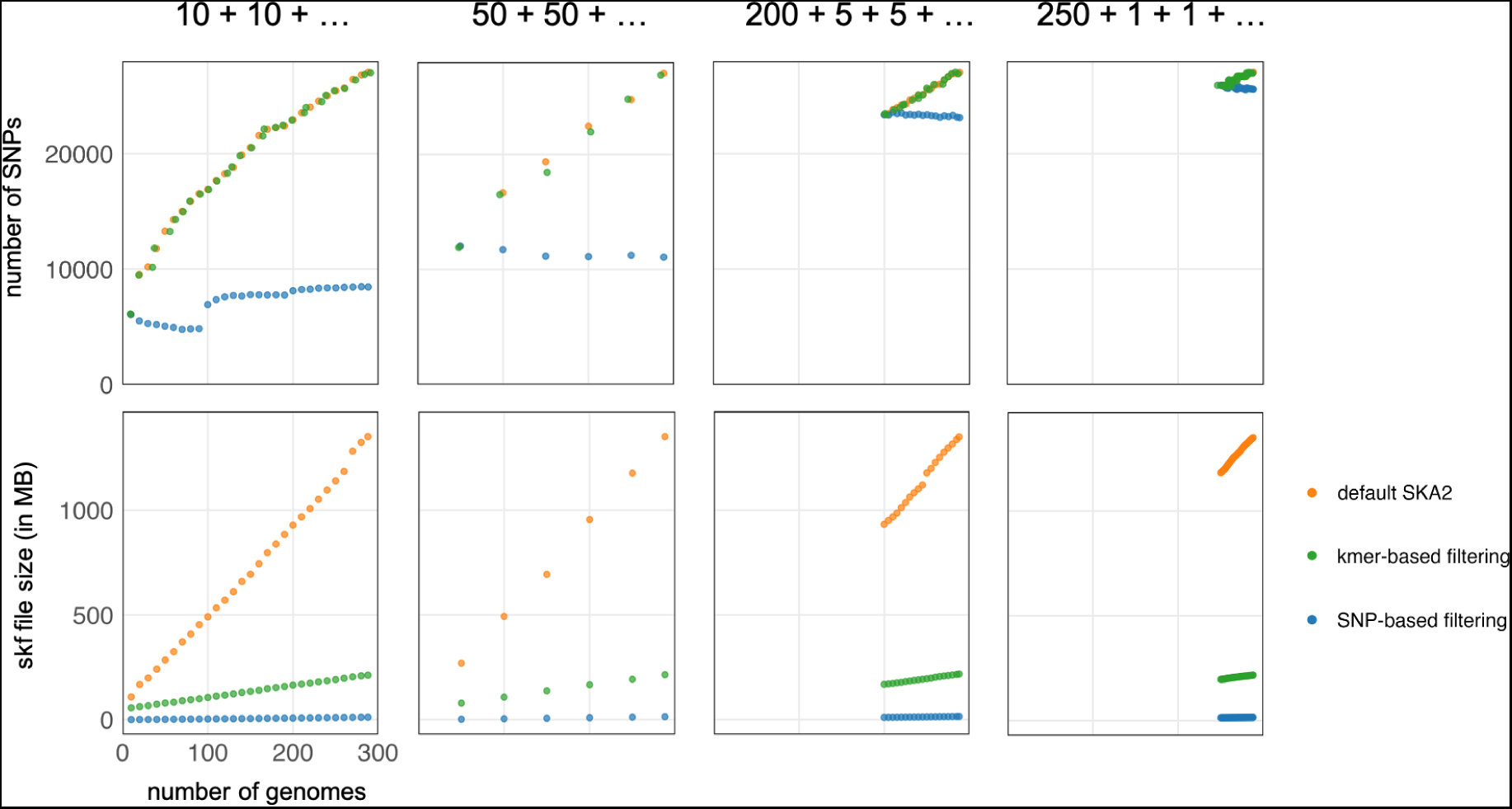
Online analyses of *E.coli* genomes. The three different genome addition strategies mentioned in the main text are displayed from left to right. Units of the x-axes (number of genomes) are identical across the six plots, and units of the y-axes (number of SNPs and skf file sizes) are identical within plots on the same line. “k-mer-based filtering” refers to the filtering based on missing split-k-mers and “SNP-based filtering” refers to the filtering based on presence-absence of SNPs. Points corresponding to the number of SNPs (upper panel) obtained from default SKA2 analyses and after kmer-based filtering were jittered to avoid a strict overlapping.

We repeated all these analyses using a stricter split k-mer filtering based on the presence-absence of SNPs, removing any constant or gap only sites (filters no-ambig-or-const and no-gap-only-sites). We observed a more substantial reduction in skf file sizes (ranging from 100-fold to 400-fold size reductions across all analyses), reaching 12-14 Mb for 288 genomes, but the numbers of missed SNPs were significant and increased in each iteration (Figure 5). Consequently, the filtering based on SNPs would only be suitable for the analysis, storage and sharing of final datasets, for example fixed SNP-based typing schemes (Hawkey et al. 2021).

## Discussion

Many popular tools in bioinformatics succeed by doing something relatively simple, but doing it well. SKA2 uses exact matching of k-mers, which has been the focus of intense optimisation efforts in bioinformatics, to create SNP alignments without explicitly performing any alignment. This is very similar to the pseudo-alignment approach which has become the dominant form of analysis of RNA-seq data (Bray et al. 2016) and for representing population data (Holley and Melsted 2020; Alanko et al. 2023; Břinda et al. 2023). SKA2 has very similar benefits – speed, ease of use, robustness to structural variation, and reduced reference bias.

In analyses of simulated outbreaks, where samples were closely related and thus most suited for split k-mer analysis, SKA2 showed higher sensitivity than mapping approaches that used distant strains as reference genomes. Another major advantage of SKA2 over classical alignment-based methods was that it avoided reference-bias or sensitivity to parameters, generating no false positives. SKA2 was also faster than other methods, with runtimes more than an order of magnitude lower than those of read-alignment pipelines and representing at least a two-fold improvement over SKA1. We further showed that iterative filtering of split k-mers represents a promising approach to track variation when streaming input samples, a potential solution for storing and analysing the ever-growing amount of publicly available bacterial genome data.

The high accuracy and speed of SKA2 are complemented by its user-friendly workflow, requiring only two commands for standard operation - one for constructing and the other for aligning split-k-mers. Additionally, akin to other alignment-free methods, outbreak investigations conducted using SKA2 eliminate the need for analysis steps such as manually masking low-complexity genomic regions (e.g. PE-PPE genes in *M. tuberculosis*) and expert selection of closely related reference genomes. Another practical advantage of SKA2 is the ability to directly and efficiently work with assembly rather than read data, or even combine the two. As sequence assemblies are becoming an increasingly popular archival form for bacterial genomes (Blackwell et al. 2021; Břinda et al. 2023; Hunt et al. 2024) they are more convenient for many users.

Algorithm scalability is of increasing importance to users. With hundreds of thousands of potential comparator genomes in the public domain (Hunt et al. 2024), users cannot afford to ‘just wait’ for computation to complete, and many who would benefit from pathogen genomics do not have easy access to high-performance computing servers. This is also important for public health pathogen genomics, where quick turn-around-time is essential to allow genomic data to be included in rapid public health decision making. Fast, simple methods that can run on regular laptops are particularly important in resource-limited settings where there is a lack of compute and skilled bioinformaticians to allow local genomic epidemiology in countries where most infectious disease outbreaks and burden are concentrated.

There is a clear potential to further develop SKA2 to accommodate emerging sequencing technologies. While we did not test SKA2’s performance using sequencing reads with high error rates (e.g. Oxford Nanopore sequencing data), we would expect the current SKA2 error filtering to perform relatively poorly on such noisy data. However, assembling this data for use as input should still be suitable (Sanderson et al. 2023), and it is also expected that advances in this technology mean that high error rate reads will likely only be an issue in the short term. We also only looked at data from single colony picks, but the identification and analysis of samples composed of a mixture of strains from plate sweeps (Mäklin et al. 2020; Mäklin et al. 2021) or metagenomic samples are becoming more common (Richardson et al. 2023). SKA2 currently only saves the observed middle-bases of each split k-mer, but a future version could also save their abundances, information which can potentially be used to deconvolute such mixtures. Also emerging are metagenome-assembled genomes (MAGs), which are typically more incomplete and contaminated than colony pick-generated assemblies (Bickhart et al. 2022).

SKA2 remains under active development. Among future improvements, better compression of the output files in SKA2 is possible, which will be important when tracking variation in very large strain collections. Future versions of SKA2 will use a sparse and phylogenetically compressed (Břinda et al. 2023) data structure for the variant array. With many constant sites this would be expected to yield an improvement, and be usable during construction too, saving memory and disk usage, and making parallelisation more effective. Various data structures for compressing k-mers also exist, and could be applied to the lists of front and back halves of the split k-mers (Rahman et al. 2021; Břinda et al. 2023). Runtimes of analyses based on sequencing reads could also be further reduced by limiting the number of reads to be analysed using a stopping criterion as implemented in other tools (Peterlongo et al. 2017; Derelle et al. 2023) and with a cache-optimised filter (Holley and Melsted 2020). Finally, the inference of variants by SKA2 is limited to SNPs, and indels or structural variants cannot be detected using the current implementation. Such variants could be inferred by repeating the split k-mer analysis at various split sizes and matching flanking bases across a middle region of various lengths, similar to haplotyping approaches (Garrison and Marth 2012), or by using split k-mers to create a de Bruijn graph as employed in other alignment-free variant calling methods (Iqbal et al. 2013; Fang et al. 2016; Peterlongo et al. 2017).

We originally released SKA in 2018, and SKA2 in 2022, so we have had time to work with users to improve testing, address unexpected results, and identify edge cases. Creating reliable results with well-chosen defaults, offering the right amount of flexibility in the interface, good documentation and long-term maintenance potential are also important to users – an important focus of our work which has only been touched upon in this paper. This has culminated in a high-quality implementation which we are confident will continue to deliver accurate results as genomic sequencing of bacteria continues to grow.

### Data and Resource Availability

The code for SKA2 is available at https://github.com/bacpop/ska.rust. Analysis code used to generate the figures in the paper is available in Supplementary Materials and at https://github.com/bacpop/ska_simulations and https://github.com/rderelle/compareALI.

## Acknowledgements

J.v.W., T. M., J. H. and J.A.L. were supported by the European Molecular Biology Laboratory. T.M. was also supported by the Academy of Finland EuroHPC grant. R.D. was supported by the NIHR Health Protection Research Unit in Respiratory Infections, in partnership with the UK Health Security Agency [NIHR200927]. N.J.C. was supported by the UK Medical Research Council and Department for International Development (grants MR/R015600/1 and MR/T016434/1). L.C. and N. J. C. acknowledge funding from the MRC Centre for Global Infectious Disease Analysis (reference MR/X020258/1 and MR/T016434/1), funded by the UK Medical Research Council (MRC). This UK funded award is carried out in the frame of the Global Health EDCTP3 Joint Undertaking.

