## Supplementary material for "Seamless, rapid and accurate analyses of outbreak genomic data using Split K-mer Analysis (SKA)"

**Supplementary figure 1:** Example output of the coverage model `ska cov` fitted to k-mer count data from Illumina sequencing.

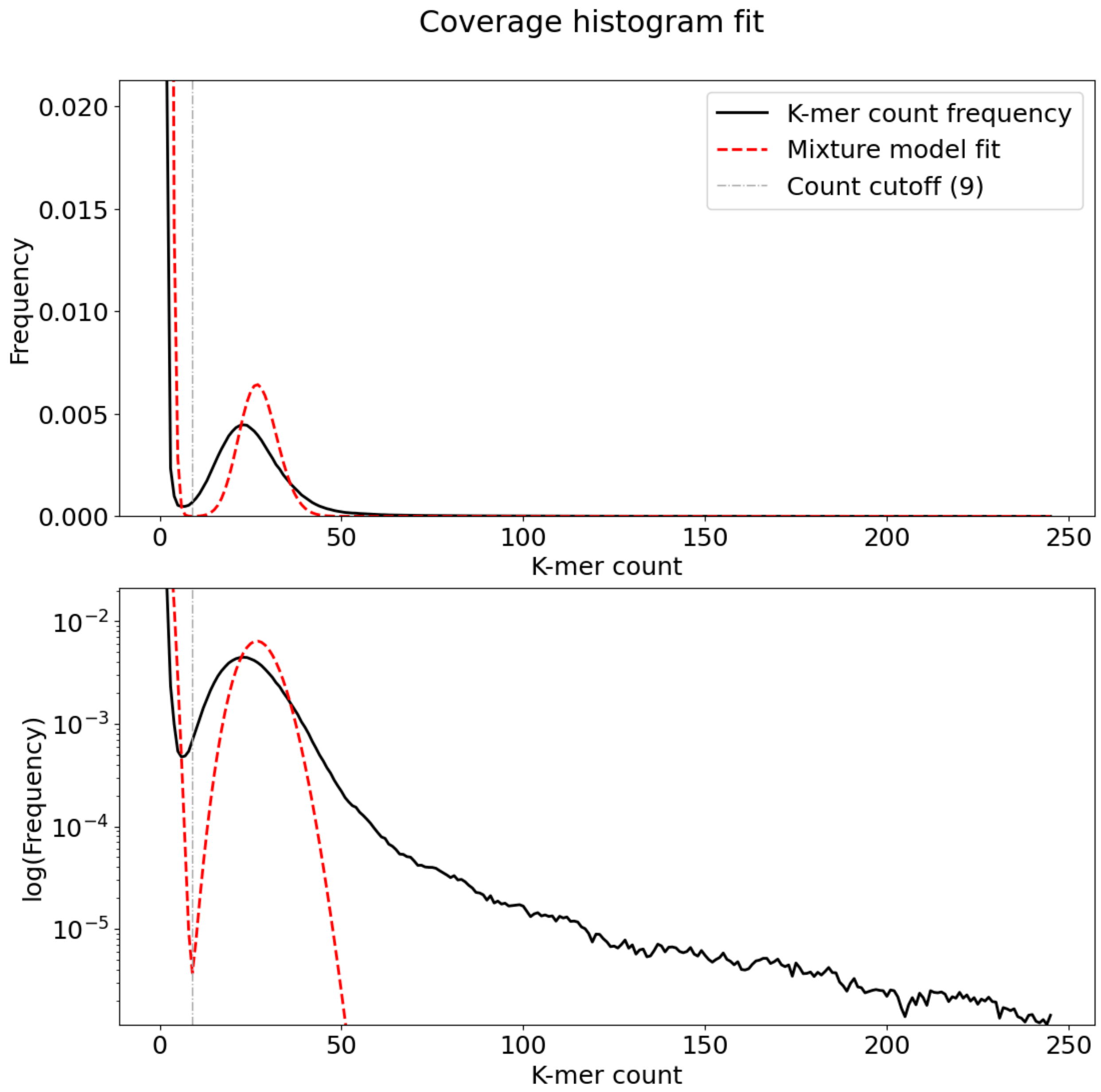

**Supplementary figure 2:** Example output of `ska distance`, connecting clusters below a given SNP threshold. Also available online https://microreact.org/project/icypNiESu31YhN1V8xzqX1-ska-distance-run-on-2023-jun-18-1730 and https://microreact.org/project/rNPR9KCnEWidVwvDmfbgQP-ska-distance-run-on-2023-jun-19-1119

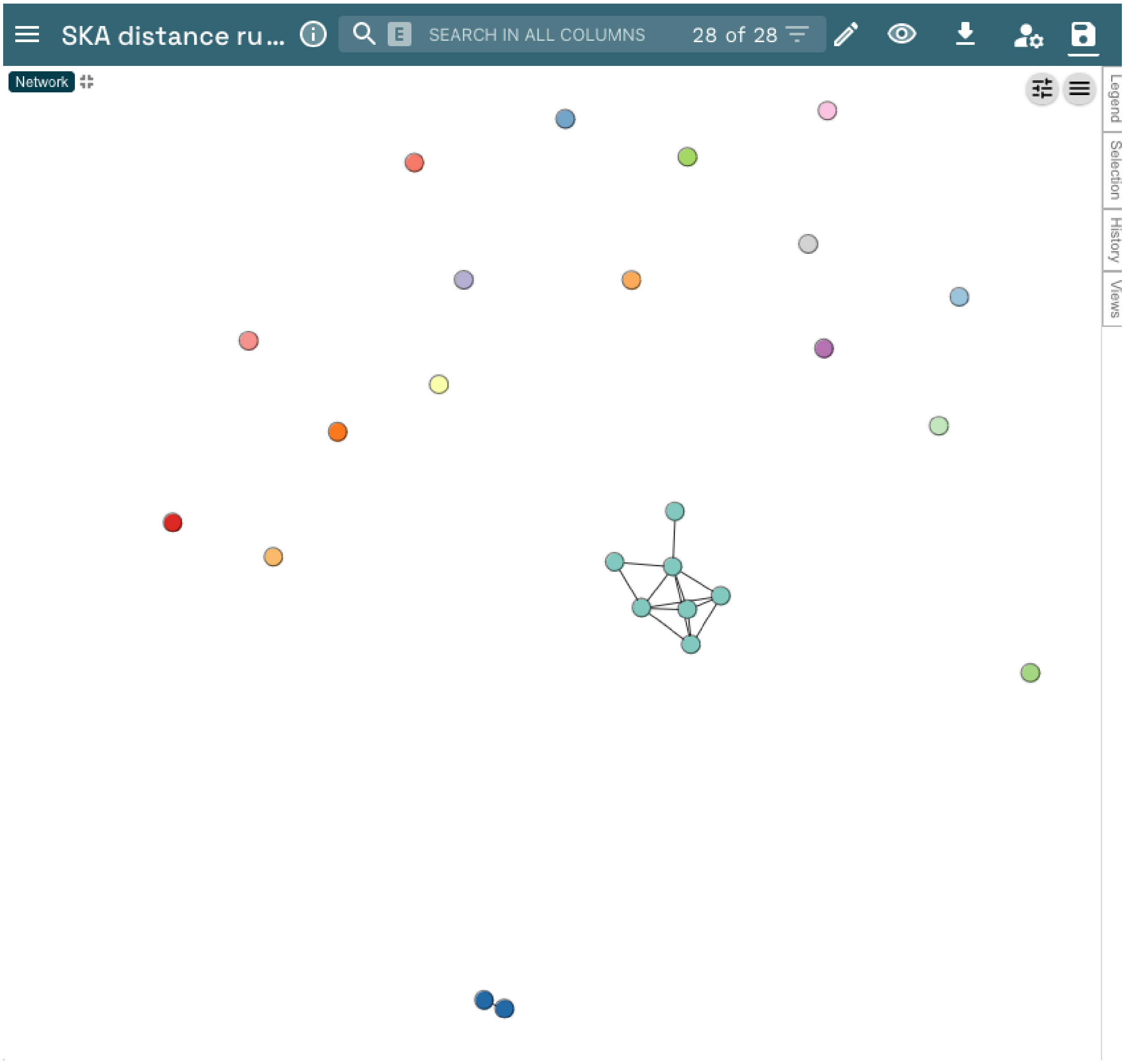

**Supplementary figure 3:** Effect of indels on SKA2's recall, same simulation setup as figure 2 but varying indel rate shown. Error bars are the 95% range from 20 repeats of the simulation.

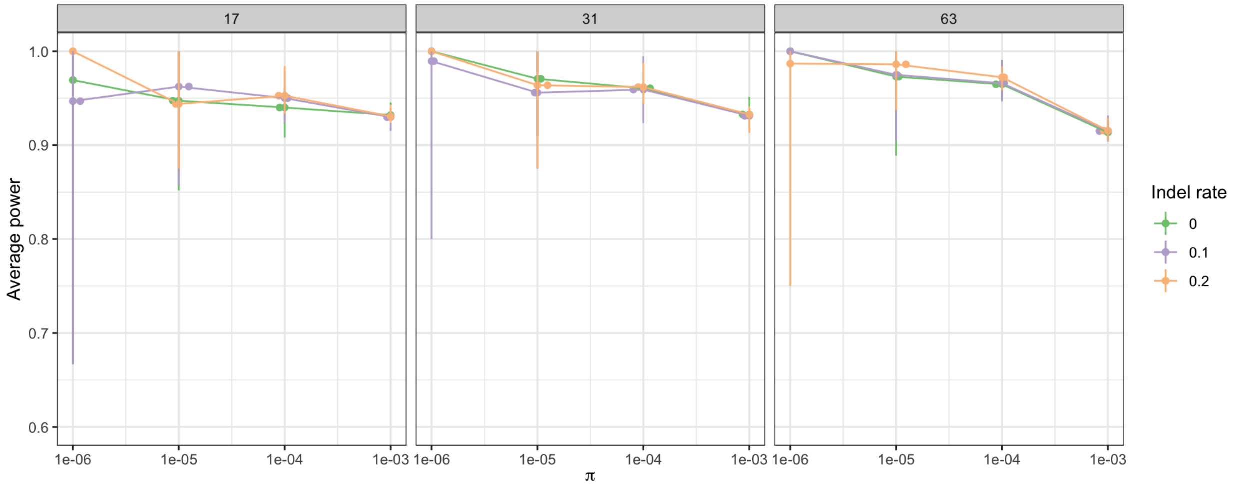

**Supplementary figure 4:** Number of SNP identified by SKA2 at different kmer sizes. The red dotted lines indicate the expected number of SNPs for each simulated outbreak.

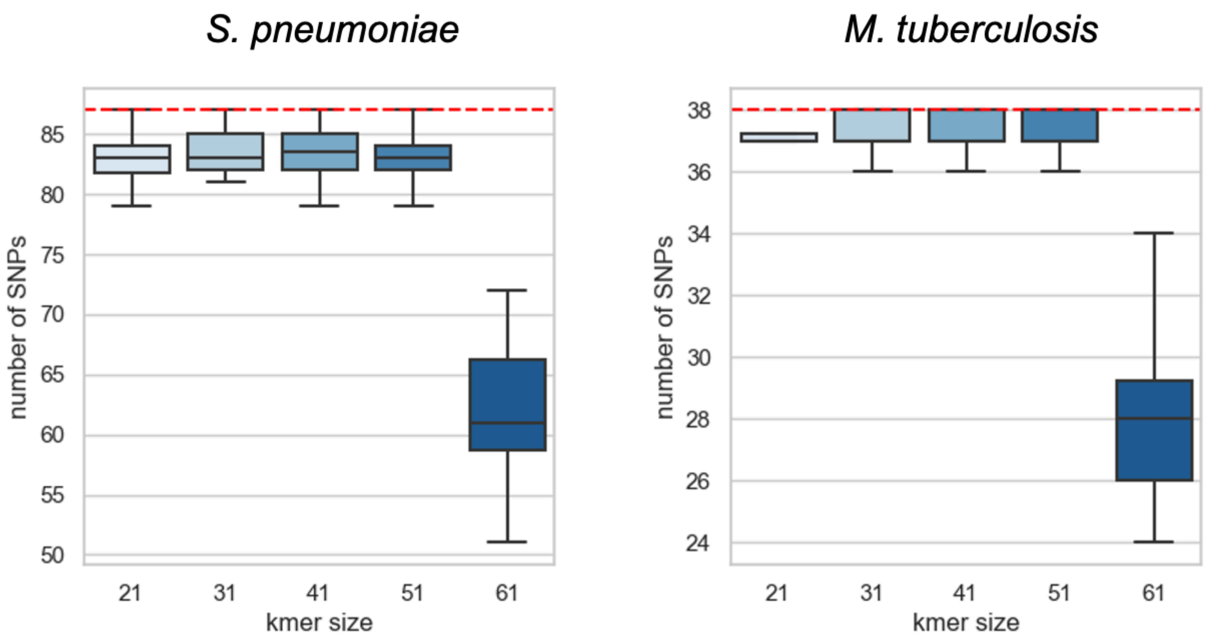

**Supplementary figure 5:** SNP detection at low coverages. The simulated outbreak analyses were repeated at low coverage settings for outbreaks generated from the strain genomes ATCC_700669 and D39V, and H37Rv and lin_4.8 for *S. pneumoniae* and *M. tuberculosis* respectively (i.e., a total of 10 simulated outbreaks at each coverage for each species). Error bars represent the 95% confidence interval.

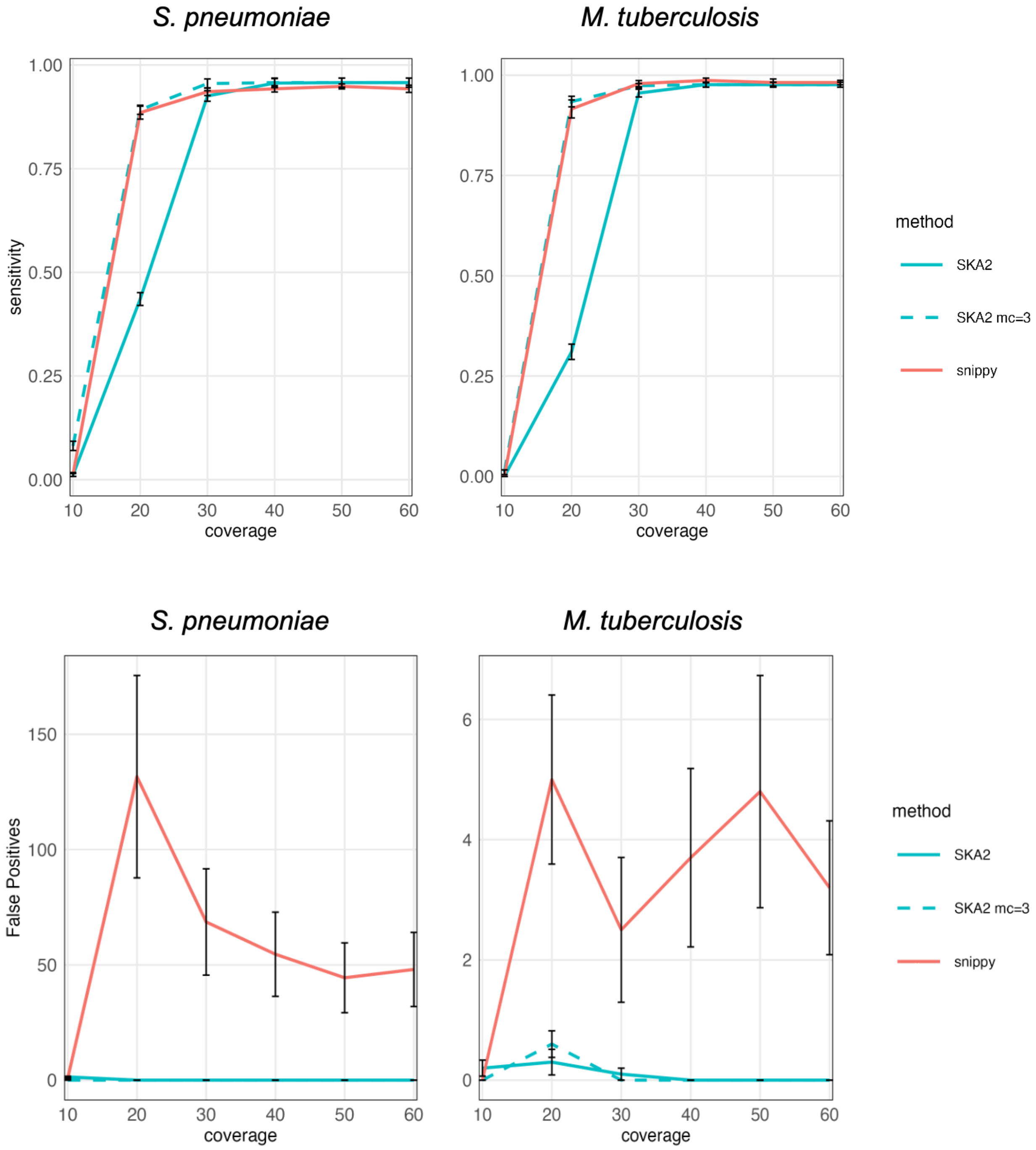

**Supplementary figure 6:** Gubbins analysis of PMEN1 samples. Top: original analysis using mapping and SNP calling from sequence reads against the Spn23F reference. Bottom: analysis using ska map. Visualised in phandango.

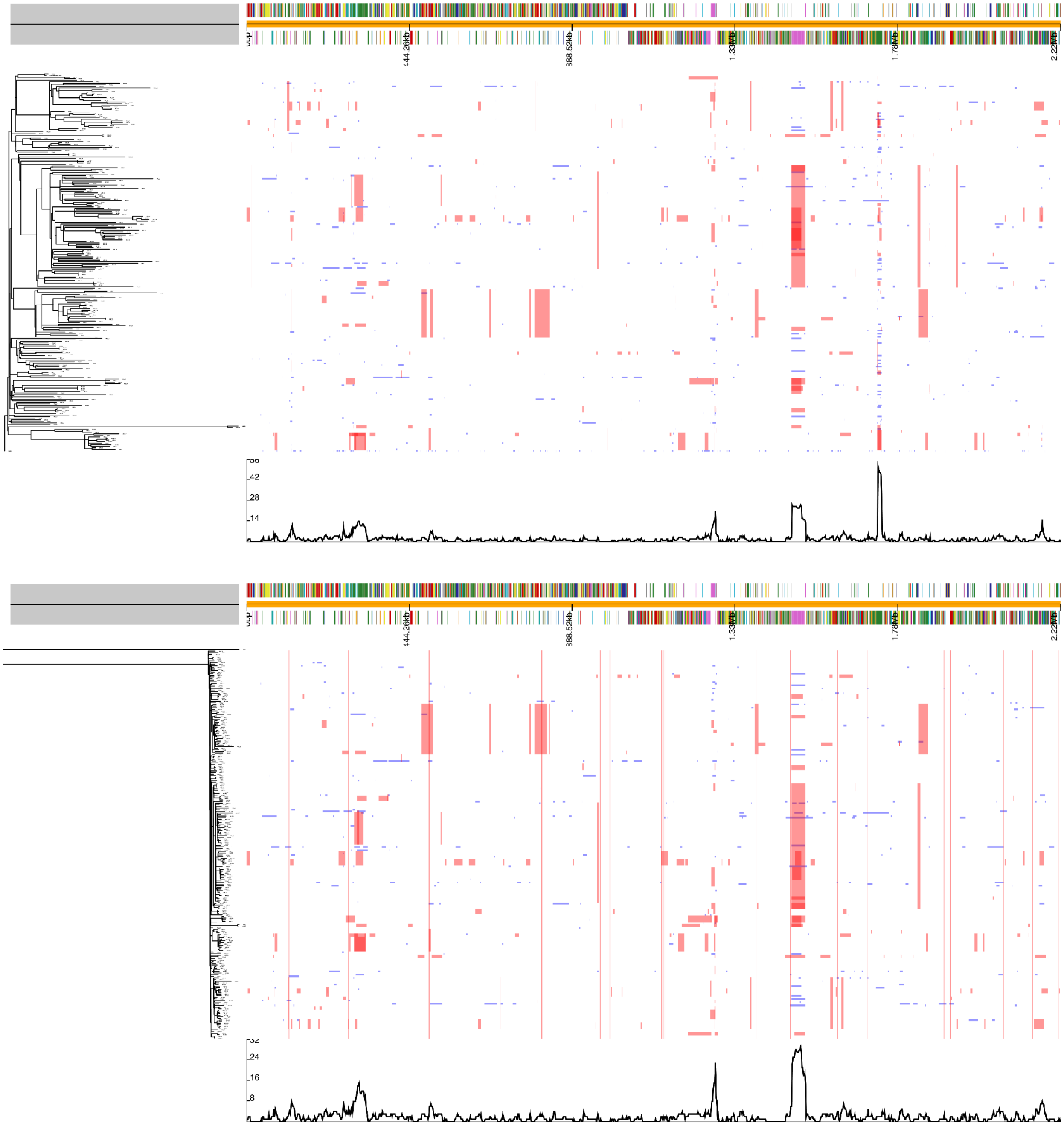

**Supplementary table 1:** Genome assembly accessions used as root and reference for the simulations of outbreaks

| strain | assembly ID |
|---|---|
| ATCC_700669 | NC_011900.1 |
| D39V | NZ_CP027540.1 |
| GPSC47 | NZ_LR216060.1 |
| Taiwan19F-14 | NC_012469.1 |

| lineage | assembly ID |
|---|---|
| H37Rv | NC_000962.3 |
| 4.8 | NZ_CP041804.1 |
| 3 | NZ_CP041871.1 |
| 1 | NZ_AP018033.1 |

**Supplementary table 2:** List of the 288 *E.coli* genome GenBank accession numbers
GCA_010806885.1 GCA_902710505.2 GCA_014482275.1 GCA_018156485.1 GCA_009684105.1
GCA_014510545.1 GCA_012714485.1 GCA_014094315.1 GCA_013026005.1 GCA_000459175.1
GCA_012351795.1 GCA_002886425.1 GCA_016655085.1 GCA_000352825.1 GCA_003308975.1
GCA_017716315.1 GCA_012856475.1 GCA_013392695.1 GCA_002466895.1 GCA_018185135.1
GCA_014137255.1 GCA_003591475.1 GCA_014643235.1 GCA_014487175.1 GCA_000459795.1
GCA_000777995.1 GCA_012376145.2 GCA_012992085.1 GCA_902668645.1 GCA_012903805.1
GCA_017004115.1 GCA_013969625.1 GCA_009683345.1 GCA_014682415.1 GCA_018194835.1
GCA_013352465.1 GCA_012788855.1 GCA_014748965.1 GCA_017075295.1 GCA_012983015.1
GCA_012297425.1 GCA_012862355.1 GCA_015208645.1 GCA_000457935.1 GCA_017066715.1
GCA_905133415.1 GCA_000714995.1 GCA_000408385.1 GCA_014477995.1 GCA_000350625.1
GCA_013181175.1 GCA_017736995.1 GCA_018173815.1 GCA_014154705.1 GCA_012952605.1
GCA_013176125.1 GCA_015620605.1 GCA_014327635.1 GCA_017132035.1 GCA_902161155.1
GCA_003145805.1 GCA_011318155.1 GCA_000503435.1 GCA_017946865.1 GCA_006230795.1
GCA_016389345.1 GCA_017747605.1 GCA_012859935.1 GCA_012561205.1 GCA_014080435.1
GCA_902859445.1 GCA_000780295.1 GCA_014477475.1 GCA_016638765.1 GCA_018170555.1
GCA_012617845.1 GCA_903978025.1 GCA_016113925.1 GCA_013024015.1 GCA_017673105.1
GCA_016786075.1 GCA_902707515.2 GCA_000776335.1 GCA_012849635.1 GCA_017722635.1
GCA_003338215.1 GCA_015195705.1 GCA_014747345.1 GCA_012694085.1 GCA_014083105.1
GCA_012766975.1 GCA_013024525.1 GCA_902841735.1 GCA_005388865.1 GCA_017841955.1
GCA_015283625.1 GCA_013076465.1 GCA_013022625.1 GCA_013964425.1 GCA_013016005.1
GCA_012875595.1 GCA_017980015.1 GCA_902827305.1 GCA_013182875.1 GCA_012366395.1
GCA_008041205.1 GCA_013008035.1 GCA_017171235.1 GCA_014140465.1 GCA_013964495.1
GCA_014143535.1 GCA_012539615.1 GCA_012171075.1 GCA_014140875.1 GCA_014140755.1
GCA_016092565.1 GCA_003773645.1 GCA_014080705.1 GCA_006234075.1 GCA_013182845.1
GCA_017065575.1 GCA_014657315.1 GCA_017066555.1 GCA_013356645.1 GCA_014081065.1
GCA_014761665.1 GCA_013028765.1 GCA_014159555.1 GCA_017679305.1 GCA_012856455.1
GCA_013022465.1 GCA_013007695.1 GCA_017737175.1 GCA_011877805.1 GCA_000776285.1
GCA_014144135.1 GCA_014099175.1 GCA_001621545.1 GCA_018173595.1 GCA_014642575.1
GCA_017064595.1 GCA_011930115.1 GCA_013964545.1 GCA_013354395.1 GCA_013041445.1
GCA_012300705.1 GCA_012865435.1 GCA_014098595.1 GCA_017721395.1 GCA_013351065.1
GCA_012162085.1 GCA_012776075.1 GCA_017747845.1 GCA_017947205.1 GCA_013017745.1
GCA_012681385.1 GCA_900500145.1 GCA_013172405.1 GCA_012752215.1 GCA_017736035.1
GCA_013129415.1 GCA_017822795.1 GCA_902708585.2 GCA_017005835.1 GCA_012864875.1
GCA_017747445.1 GCA_013026915.1 GCA_014749705.1 GCA_012478295.1 GCA_013072125.1

GCA_002002255.1 GCA_009683215.1 GCA_002109565.1 GCA_017737035.1 GCA_000459815.1 GCA_018167175.1 GCA_013968525.1 GCA_902710295.2 GCA_009729815.1 GCA_015042015.1 GCA_015163115.1 GCA_017673085.1 GCA_014772575.1 GCA_014902335.1 GCA_902707215.2 GCA_012876505.1 GCA_013512695.1 GCA_012384675.2 GCA_016242675.1 GCA_000458515.1 GCA_014429545.1 GCA_014470215.1 GCA_012972545.1 GCA_012601635.1 GCA_012453665.1 GCA_003795545.1 GCA_902849475.1 GCA_013083485.1 GCA_014421945.1 GCA_018167975.1 GCA_012169475.1 GCA_003322335.1 GCA_009790015.1 GCA_012680905.1 GCA_012603555.1 GCA_015644435.1 GCA_017822615.1 GCA_012260985.1 GCA_016574135.1 GCA_902709295.2 GCA_902840725.1 GCA_017779405.1 GCA_012599375.1 GCA_014099615.1 GCA_012672805.1 GCA_018164155.1 GCA_015138385.1 GCA_012296825.1 GCA_013062025.1 GCA_012008175.1 GCA_017001175.1 GCA_012700325.1 GCA_012117835.1 GCA_013068565.1 GCA_009790475.1 GCA_902708505.2 GCA_014463365.1 GCA_905133355.1 GCA_014082825.1 GCA_014935385.1 GCA_014733035.1 GCA_012871915.1 GCA_001749565.1 GCA_015208465.1 GCA_000326945.1 GCA_012330165.1 GCA_012870815.1 GCA_014098395.1 GCA_000026325.2 GCA_902709885.2 GCA_000456925.1 GCA_902708615.2 GCA_017736055.1 GCA_903977735.1 GCA_014479615.1 GCA_012564745.1 GCA_012756775.1 GCA_017002755.1 GCA_012774335.1 GCA_012474695.1 GCA_012642785.1 GCA_000418635.1 GCA_013972545.1 GCA_014775885.1 GCA_012225765.1 GCA_013008135.1 GCA_012434655.1 GCA_014779295.1 GCA_015288625.1 GCA_017787665.1 GCA_003892475.1 GCA_012873715.1 GCA_902838585.1 GCA_000711415.1 GCA_902847965.1 GCA_012695305.1 GCA_012862105.1 GCA_902710035.2 GCA_900480125.1 GCA_012759815.1 GCA_013009735.1 GCA_008041395.1 GCA_000458725.1 GCA_012910105.1 GCA_012805155.1 GCA_009766465.1 GCA_014082595.1 GCA_017034675.1 GCA_012356355.1 GCA_000164195.1 GCA_009683405.1 GCA_017052855.1 GCA_012871755.1 GCA_009680755.1 GCA_017066615.1 GCA_003885225.1 GCA_012670565.1 GCA_904419595.1

**Supplementary data 1:** Details of parameters used in the outbreak simulation runs

<u>Transphylo parameters:</u>

| | *S. pneumoniae* | *M. tuberculosis* |
|---|---|---|
| Neg | 250/365 | 100/365 |
| w.scale | 1.5625 | 0.1 |
| w.shape | 1 | 10 |
| pi | 0.5 | 0.25 |
| off.r | 1.5 | 5 |
| dateStartOutbreak | 2005 | 2005 |
| dateT | 2007 | 2009 |
| nSampled | 12 | 30 |

<u>phastSim command lines</u>

_ *M. tuberculosis*:

phastSim --outpath test_sim --mutationRates HKY85 0.23 0.5 0.17 0.33 0.33 0.17 --reference H37Rv.fna --treeFile mod_TransPhylo.tre --seed 0 --rootGenomeFrequencies 0

_ *S. pneumoniae*:

phastSim --outpath test_sim --mutationRates HKY85 0.23 0.5 0.17 0.33 0.33 0.17 --reference Spn_ATCC_700669.fna --treeFile mod_TransPhylo.tre --seed 0 --insertionRate CONSTANT 0.0876 --deletionRate CONSTANT 0.0876 --insertionLength DISCRETE 0 0 0 0 0 0 0 0 0 0 0 0 0 0 0 0 0 0 0 0 0 0 0 0 0 0 0 0 0 0 0 0 0 0 0 0 0 0 0 0 0 0 0 0 0 0 0 0 0 0 0 0 0 0 0 0 0 0 0 0 0 0 0 0 0 0 0 0 0 0 0 0 0 0 0 0 0 0 0 0 0 0 0 0 0 0 0 0 0 0 0 0 0 0 0 0 0 0 0 1 --deletionLength DISCRETE 0 0 0 0 0 0 0 0 0 0 0 0 0 0 0 0 0 0 0 0 0 0 0 0 0 0 0 0 0 0 0 0 0 0 0 0 0 0 0 0 0 0 0 0 0 0 0 0 0 0 0 0 0 0 0 0 0 0 0 0 0 0 0 0 0 0 0 0 0 0 0 0 0 0 0 0 0 0 0 0 0 0 0 0 0 0 0 0 0 0 0 0 0 0 0 0 0 0 0 1 --indels --rootGenomeFrequencies 0

<u>Command-lines of the BWA+BCFtools custom pipeline:</u>

bwa mem -t 1 -o sample.sam reference_genome.fna sample_1.fq.gz sample_2.fq.gz
samtools view -q 20 -bS sample.sam > sample.bam
samtools sort -m 8G -o sample.sorted.bam sample.bam
samtools depth -Q 20 -q 20 -a sample.sorted.bam > sample_cov.txt
bcftools mpileup -Q 20 -f Spn_ATCC_700669.fna sample.sorted.bam -o sample.vcf1
bcftools call -f GQ -o sample_SNPs.vcf -V indels -cv sample.vcf1
